# Targeted downregulation of estradiol binding Na^+^/H^+^ exchanger *nhx-2,* mimics calorie restriction, extends reproductive longevity and ameliorates effects associated with alpha synuclein aggregation in *C. elegans*

**DOI:** 10.1101/2020.07.30.229344

**Authors:** Shikha Shukla, Lalit Kumar, Arunabh Sarkar, Kottapalli Srividya, Aamir Nazir

## Abstract

Setting in of reproductive senescence (RS) gives rise to several changes, making aged individuals susceptible to multiple disorders including neurodegenerative diseases, cardiovascular ailments and bone disorders amongst others. The present study, employing transgenic *C. elegans* that expresses ‘human’ alpha synuclein, endeavors to decipher the association of reproductive senescence with age-associated neurodegenerative diseases and behavioral ageing, under normal conditions and after being probed with estradiol. We carried out RNAi induced silencing of a subset of 22 genes that are known to delay RS, followed by studies on alpha-Synuclein aggregation and associated effects. These studies led us to functional characterization of the Na^+^/H^+^ exchanger; *nhx-2*, expressed exclusively in gut. We found that RNAi of *nhx-2* not only ameliorates the effects associated with alpha-Synuclein aggregation, but it also attunes effects related to behavioral aging including that of reproductive health-span and neuroprotection via mimicking dietary restriction, as it alters food absorption from the gut. We further elucidated that these effects are Sir-2.1 driven as *nhx-2* knock out did not delay reproductive senescence in knock down condition of *sir-2.1.* To substantiate our findings, we performed whole transcriptome analysis in *nhx-2* mutant strain. Our data revealed differential expression of 61 out of 62 hallmark genes of CR described by GenDR, in knock out condition of *nhx-2.* As estradiol plays a central role in both reproductive health as well as neuronal health, we subjected worms to exogenous estradiol treatment and observed that it led to elevated levels of *nhx-2*. Studies on structural binding analysis demonstrated significant binding potential of estradiol receptor NHR-14 with *nhx-2* gene and ChIP analysis revealed that estradiol treatment gives rise to enhanced NHX-2 levels through inducing the promoter specific histone H3 acetylation (H3K9) and lysine methylation (H3K4me3). These studies identify *nhx-2* as an important modulator that extends reproductive longevity and ameliorates effects associated with alpha synuclein aggregation in *C elegans*.

**Graphical abstract:** 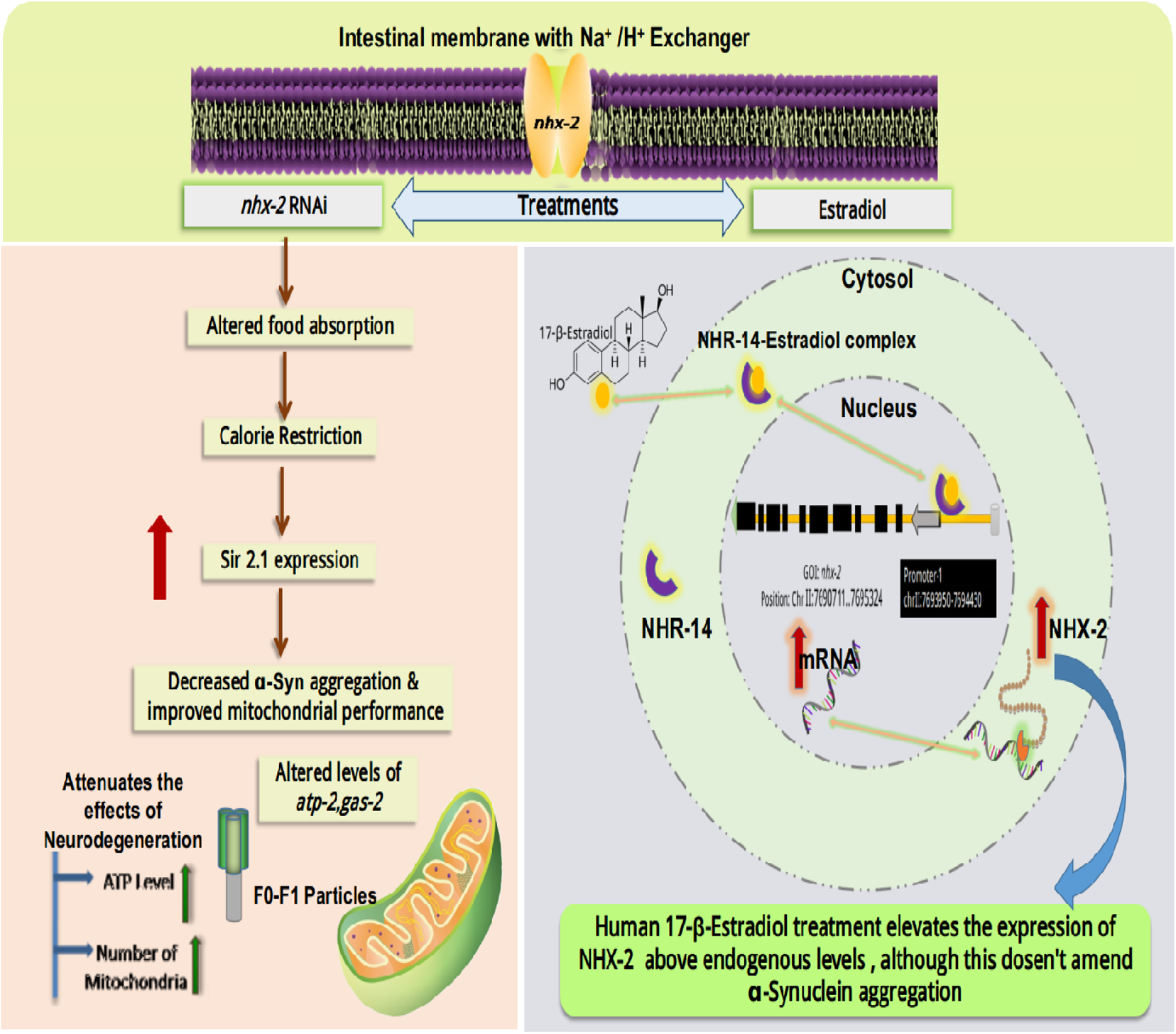

**HIGHLIGHTS:**

1. Silencing of sodium proton antiporter *nhx-2* affords neuroprotection and ameliorates effects associated with alpha-Synuclein aggregation via mimicking dietary restriction in *C. elegans*.
2. Exogenous 17-β-Estradiol treatment induces the expression of *nhx-2,* through inducing the promoter specific histone H3 acetylation (H3K9) and lysine methylation (H3K4me3).
3. Effects associated with *nhx-2,* including prolonged reproductive span and neuroprotective effects, are SIR-2.1 driven.
4. *nhx-2* silencing decreases alpha-Synuclein aggregation, however; estradiol mediated overexpression above the endogenous levels, does not amend the aggregation any further.

## Introduction

Neurodegenerative diseases (NDs) are multifactorial ailments involving various pathological mechanisms. The most relevant and known risk factors for neurodegenerative diseases include age, familial occurrence, traumatic brain injury and cardiovascular diseases [1]. There are other potential factors that add on to the susceptibility of the disease including, inflammation, oxidative stress, hypertension, gender and endocrine imbalance amongst others [2]. The major challenge in treating NDs is; their multifactorial nature, and lack of complete understanding of associated etiology [3]. This may be the reason why most age-associated NDs have no cure yet. Hence, it has become imperative to delve deeper into the understanding of multiple processes associated with ageing which may aid in devising complete cure for these ailments. One of the hallmark features of aging is “reproductive senescence” (RS) - a gradual process that starts from the age of 35 and reaches its culmination at around 51 years of age [4]. RS is characterized by several changes including a considerable change in the levels of sex steroid hormones which happen to be potent regulators of neuronal survival [5] [6] [7] [8]. RS is not only followed by vasomotor symptoms like hot flushes, irritability, insomnia and mood disability, but it may also make a person more vulnerable to several serious health conditions. For instance, there are reports that it might put one at higher risk of osteoporosis, heart diseases, diabetes, hypertension, and breast cancer [9], thus making its association with NDs coherent. A study has shown, a low to moderate level of anxiety, depression, social dysfunction, and somatic symptoms and psychosocial stress in women going through reproductive senescence [10] [11]. Biochemical and neurophysiological studies report that fluctuations in these hormones, followed by reproductive senescence may affect cognition. It may also affect the cholinergic and serotonergic activity in specific regions of the brain [12], maintenance of neural circuitry, favorable lipoprotein alterations, and prevention of cerebral ischemia [13]. Experimental, epidemiological and clinical evidence suggests, that sex steroid hormones have neuroprotective actions. Levels of estrogen and progesterone play a very critical role in the development of neurodegenerative diseases [14]. This perspective of pathology advocates that a variety of neurodegenerative diseases can be caused by reproductive senescence which causes a decline in sex steroid level and provides a novel therapeutic approach that may affect a broad spectrum of neurodegenerative diseases. Taking clues from these facts, we hypothesize that molecular changes associated with reproductive senescence and gene expression modulations; induced by altered hormonal levels might be associated with neurodegenerative diseases.

*C. elegans* has been employed to study ageing and reproductive framework with several genetic approaches as well as genome wide screenings. There are various molecular pathways that are extended over in *C. elegans* and humans. *C. elegans* discontinue laying eggs after one third of its lifespan and the chromosomal aberrations for instance; chromosome non-disjunction is one of the hallmark features of germ line ageing [15]. These aberrant molecular events are very well demonstrated in humans and signifies that reproductive senescence holds common molecular changes in *C. elegans* as well as humans. In the present study, we have carried out an in-depth analysis on RS related gene and its association with neurodegeneration having an aim of deciphering the link between healthy aging/longevity induced via prolonged reproductive health span and its impact on neuronal health in *C. elegans*. We studied gene that showed an affinity with the estradiol receptor, and whose expressions are modulated upon the treatment of estradiol as this hormone is known to play a central role in reproductive senescence and neuroprotection. The course of our studies vindicated the interplay of reproductive span, gut homeostasis, calorie restriction and neuroprotection via sirtuins. Herein, we found an interestingly overlapping RS, ND and gut health under an umbrella and elucidated that Sir-2.1 is the central ground of the underlying mechanisms involved in neuroprotection.

Our study, elucidates the association between RS, gut health, estradiol induced modulation of genes and their association with neurodegenerative disease.

## Methods

### Strains used

***Bristol strain N2*** (wild type)

***NL5901*** (Punc-54::alpha-Synuclein:: YFP + unc-119)

***VC199*** (ok434) *sir-2.1*

***UL2992*** [*sir-2.1*::mCherry + rol-6(su1006)]

***VC363*** (ok553)/mIn1 [mIs14dpy-10(e128)] II

***BZ555*** [dat-1p::GFP]

***AN105*** [sir-2.1::mCherry + rol-6(su1006::Punc-54::alpha-Synuclein:: YFP + unc-119] This study

### *C. elegans* strains and maintenance

Bristol strain N2 and NL5901 strain expressing “human” alpha-Synuclein protein in the muscles with YFP expression, VC199, UL2992 expressing *sir-2.1* ubiquitously, VC363 (*nhx-2* knock out) obtained from the Caenorhabditis Genetic Center, University of Minnesota, St. Paul, MN, USA) were used in this study. *C. elegans* strains were cultured using standard protocols as described previously [16]. *Escherichia coli (E.coli)* strain OP50 was used as standard food to feed the worms. Synchronized worms were obtained by embryo isolation procedure as described [17]. Briefly, worms were washed with M9 buffer and treated with axenizing solution (2 ml of sodium hypochlorite and 5 ml of 1 M sodium hydroxide solution) until eggs were released from the body.

### RNAi-induced gene silencing

Gene silencing was carried out as described previously employing the feeding protocol for the silencing of genes of interest [18]. The dsRNA targeted for *C. elegans* gene expressed in bacterial clone (source Ahringer library) was cultured for 6–8 h in LB containing ampicillin (50 μg/ml). This bacterial culture was seeded onto NGM plates having 5 mM IPTG and 25 mg/L carbenicillin and incubated at 37 °C for 6-8 hours. Age-synchronous embryos of worms were transferred onto these plates for further experiments.

#### Fluorescent imaging of alpha-Synuclein protein and dopaminergic neurons

To analyze the effect of *nhx-2* silencing on alpha-Synuclein aggregation, around 300 worms were grown on IPTG plates seeded with specific bacterial clones expressing dsRNA targeted at *nhx-2*. After 48 hours, worms were washed with M9 buffer to remove bacteria and immobilized with 100mM sodium azide (Sigma, Cat No. 71289). The imaging of live immobilized worms was carried out using a fluorescence microscope (Carl Zeiss Axio Imager M3). alpha-Synuclein expression was quantified through ImageJ software (ImageJ, National Institutes of Health, Bethesda, MD), and for fluorescence intensity measurement, 25 worms were selected from every group to measure alpha-Synuclein expression [19]. Each experiment was repeated three times. In order to perform imaging of dopaminergic neurons we used a transgenic strain BZ555[dat-1p::GFP] that expressed GFP dopamine transporters in dopaminergic neurons and treated these worms in both EV and *nhx-2* RNAi conditions and performed confocal imaging of 25 worms per group. In order to treat worms with 6-OHDA, 10 μL of the 6-OHDA 5x stock solution was freshly prepared in dH2O and added to 30 μL of L1-stage larvae in M9 [20].

#### Quantification of Reactive Oxygen Species

Quantification of ROS was performed in *C. elegans* by using 2,7-dichlorodihydrofluorescein diacetate (H_2_DCFDA) (Invitrogen Cat. No. D399) as described previously [21]. Briefly, stock solution (10 mM) was prepared fresh in absolute alcohol, which was diluted (100 μ in phosphate buffer saline (for 1 X PBS Na_2_HPO_4_ 1.44 g, KH_2_PO_4_ 240 mg, NaCl 8 g, and KCl 200 mg). Age-synchronized wild-type (N2) worms were raised either on bacteria with empty vector (EV) or bacteria-specific for RNAi of various genes of interest. After incubation for 48 h at 22 °C, worms were harvested using M9 buffer (for 1L KH_2_PO_4_ 3 g, Na_2_HPO_4_ 6 g, NaCl 5 g, and 1MMgSO_4_ 1 ml) and suspended into PBS. Approximately 100 worms (following mean counting method) in 100 μ PBS were transferred into 96-well black plates (Perkin Elmer optiplate-96 F). H_2_DCFDA at a concentration of 50 Mm was added. Fluorescence intensity was quantified at 0 min after the addition of dye and 60 min after the addition of dye using Perkin Elmer multimode plate reader (Victor X3) at excitation 485 nm and emission of 520 nm. All the samples were estimated in triplicates, and for each group 300 worms were used.

### Thrashing assay

Thrashing assay was performed with different treatment groups and the control group (EV). Worms were washed with M9 buffer and suspended on a glass slide in a drop of M9 buffer. Thrashes were counted as the bending of the head to the outermost angle of the body and then back to the previous posture of the body [22]. Thrashing assay is considered to be a reliable behavioral assay to find out the levels of dopamine in the worms as thrashes are controlled by the dopamine concentration in *C. elegans*. 25 worms per group were counted for a mean number of thrashes per group and represented in a bar graph.

### Food absorption assay

Food absorption assay was performed to check the amount of food absorbed in control as well as in RNAi conditions of *nhx-2*. After 48 hours, age synchronized worms were fed with fluorescent OP50 and were transferred on empty NGM plates for 1-2 hours to check the level of fluorescence that remained in the gut which signifies the bacteria that are still present in the gut and have not been absorbed by the worms; control group was studied as reference [23]. Fluorescence was visualized with the help of confocal microscopy.

### *in silico* studies

a) Prediction of the last PolyA signal and extracting invert repeats from the genes of interest, gene sequences of *nhx-2* & *her-1* were obtained from Wormbase. PolyApred predicted the last PolyA signal of query gene sequences based on the score [24]. RepEX server gave inverse repeats for the same query string and length was parameterized between 10-20 bases [25]. The contigs that were located between the last polyA signal and exon of the query gene were taken as a template for further steps. b) For the preparation of 3D-DNA, the contigs obtained from the prior step were given as a query to 3D-DART server for constructing the DNA with 3D coordinates [26]. The parameters were kept as the default set of instructions by the server itself. The PDB files of the respective DNA-contigs were taken to the next step. c) Docking of β-estradiol with the NHR-14 receptor (UniProtKB - 002151). 3D structures of estradiol and NHR-14 were downloaded from PubChem and UniProt respectively. Docking site prediction was done using PrankWeb (Jendele et al., 2019). From this, active sites were taken for the docking grid. Docking was performed using AutoDock tool [27]. From the docking results, the conformer with the least energy pose was taken as a hit for further steps. d) Docking of the NHR-14-estradiol complex with three different DNA contigs from two genes (*her-2* and *nhx-2*) to identify the potential binding affinity. Hex docking platform was used for the purpose where DNA contig was taken as a receptor and NHR-14 complex with estradiol was taken as a ligand [28]. The resultant complex was then processed by CASTp for binding site analysis to visualize the binding pocket and interactive residues [29].

### Quantitative Real-Time PCR (qPCR) studies for mRNA expression of various genes

The total RNA of the worms from control and experimental groups was extracted followed by the preparation of cDNA and quantification of mRNA levels of genes, employing qPCR [30]. In brief, for RNA preparation, harvested worms were washed with DEPC (Sigma, Cat. No. D5758) treated water, and the RNA was isolated using RNAzol (Molecular Research Center, Cat. No. RN190) reagent. Isolated total RNA was then subjected to cDNA synthesis using Revert Aid Premium First Strand cDNA Synthesis Kit (Fermentas, Cat. No. K1652). The cDNA was used at a concentration of 125 ng and processed for quantitative real-time PCR using SYBR green master mix to quantitate the mRNA levels of genes, using actin as the loading control.

The amplification in 96-well plate was done with the program as (1) 1 cycle of Pre-incubation: 50°C for 2 minutes and 95°C for 10 minutes; (2) 40 cycles for Amplification: 95°C for 30 seconds, 55°C for 30 seconds, 72°C for 30 seconds; (3) Melting curve analysis: 95°C for 5 seconds, 65°C for 1 minute, (4) Cooling: 40°C for 5 minutes. After amplification, the Ct values were obtained which were used for the relative quantification of the mRNA expression of the target genes. The relative quantification was based on the 2−ΔΔCt method.

### Table: 1 – Sequences of primers used

**Table 1:**
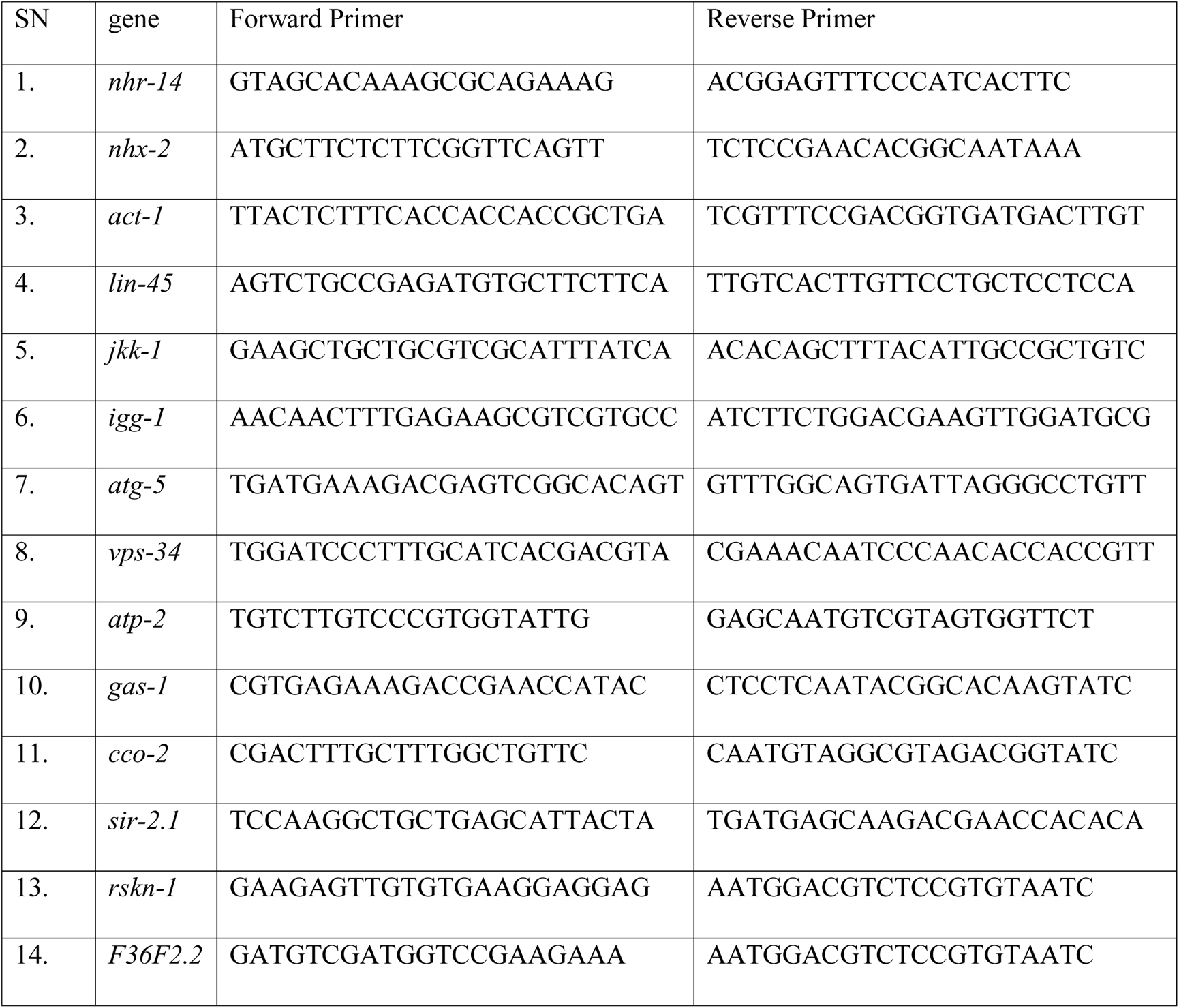
Compilation of docking analysis

#### Western blotting

In order to validate the expression studies carried out by fluorescence microscopy of GFP expressing strains, western blotting was essentially performed as described here [30]. Worms were washed thrice with M9 buffer and re-suspended in phosphate buffer saline (PBS). These worms were sonicated in PBS containing protease inhibiter cocktail (Thermo Scientific 87785). Sonication was performed at 25 amplitude for 3 min (pulse time of 15 seconds); using sonicator (Misonix S4000). The sonicated lysate was centrifuged at 16,000X g for 30 min at 4°C for removing debris and supernatant contains total protein. Protein concentration was quantified using Bradford reagent (Sigma B6916). From each group; 200µl of total protein was further fractionated into soluble and aggregated forms using membrane based centrifugal filter unit (Millipore MRCF0R100) 500X g for 15 min at 4°C. This filter allows passing of the biomolecules with molecular weight cut-off of 100 kDa. The filtrate (soluble form) and un-filtrated (aggregated form) volume was made up to total protein (200µl) using PBS. 15µg of total protein was loaded as calculated from Bradford and same amount of soluble and aggregated are loaded in 12 % sodium dodecyl sulfate polyacrylamide gel electrophoresis (SDS-PAGE). The primary antibodies used for western blot were monoclonal anti alpha-synuclein (1:1000, abcam ab138501) and anti-tubulin antibody (1:1000, abcam, ab6160). Chemiluminescence detection was performed using LAS 3000 GE ImageQuent and densitometric analysis was performed by ImageJ software (Image J, National Institute of health, Bethesda, MD).

### Fecundity assay

To perform fecundity assay ten worms (age synchronized L4 worms) of each group N2, NL5901, *nhx-2* knock out and *nhx-2* knock out in RNAi condition of *sir-2.1* were transferred on four different 90mm NGM plates and IPTG-NGM plates respectively. Worms were picked to fresh plates every day and their eggs/offspring were recorded until, there were no eggs remained on the plates in all the groups.

#### Genetic cross for creating transgenic strain that expresses mCherry Sir-2.1 and alpha-Synuclein

To create a transgenic strain that expresses mCherry SIR-2.1 ubiquitously and alpha-Synuclein intact within its muscles, we performed genetic crosses between UL2992 and NL5901 followed by backcrossing with N2 [31]. For this, standard procedures were followed, briefly, we first generated males of NL5901 strain by exposing L4 staged worms with mild temperature stress (30°c for 4-5 hours) and then placed the worm on normal 22°c which induced male formation in next generation. To perform crossing we placed 8-10 worms of NL5901 and 3-4 hermaphrodites of UL2992 on 35 mm plates. We further chose those hermaphrodites from the progeny that showed alpha-Synuclein expression in muscles. To stabilize the strain, we performed back cross with wild type. This cross generated a strain that expressed mCherry *SIR-2.1* ubiquitously and alpha-Synuclein in muscles.

### Measurement of mitochondrial content with the help of mitotracker

Mitotracker assay is an advantageous method to access mitophagy as well as mitochondrial content [32]. We employed mitotracker based mitochondrial staining to check the mitochondrial content in knock down condition of *nhx-2*. To check the effect of RNAi of *nhx-2* on mitochondrial content, worms were treated with mitotracker. To do so around, 300 worms were grown on IPTG plates having RNAi colony of strain HT115 to knock down *nhx-2*. After 48 hours, worms were washed with M9 buffer to remove bacteria and treated with Mitotracker (500 nM; Invitrogen cat. no M22426).

### ATP measurement

Adenosine 5’-triphosphate (ATP) Bioluminescent Assay Kit (Sigma FLAA) has been used for ATP detection. Standard protocol with modifications, have been utilized for ATP detection in *C. elegans*. Briefly, around 300 worms were collected from each group and proceeded for protein isolation as described above [32]. The Bradford reagent is utilized for the measurement of the concentration of protein. After that 100µl ATP assay mix (2X10-12 stock) added into control and treatment vials and incubated for 3 min at room temperature. Further 50ug in 100 µl protein sample was added into vials and for ATP measurement luminescence was measured through luminometer (Promega glowmax).

### Cell isolation and MMP measurement

#### Cell isolation

Method described here has been followed for the cell isolation experiments [33]. Age synchronized worms were pelleted via centrifugation at 13000 rpm for 2 min. Double distilled water was used for the washing of worm pellets.100 µl of the pellet was incubated for 4 min. in 200 µl of SDS-DTT solution (solution contains 0.25% SDS, 200 mM DTT, 20 mM HEPES, pH 8.0 and 3% sucrose) at room temperature. After treatment, 800 µl of egg buffer (containing 118 mM NaCl, 48 mM KCl, 2mM CaCl_2_, 2mM MgCl_2_, 25mM HEPES, pH 7.3) was added to the reaction. Worms were centrifuged at 13,000 rpm for 1 min in egg buffer and we repeated this step 5 times. For worm cuticle digestion, 15 mg/ml pronase (Sigma-Aldrich from *Streptomyces griseus*) enzyme has been used. Pronase added sample was incubated at room temperature for 20 min and during incubation for mechanical disruption we kept pipetting it. L-15 medium (Sigma-Aldrich L1518) was used for the termination of the digestion process and centrifuged at 180g for 5min on 4°C for sedimentation of cells. Cells were re-suspended in 1 ml L-15 medium and kept on ice for 15 min. 800 µl supernatant was separated into the fresh micro-centrifuge tube after ice settling step and centrifuged at 180g for 5min on 4°C

### Mitochondrial membrane potential assay

For measurement of mitochondrial membrane potential worm cells were isolated as described above sub-section. Approximately, 1X105 cells/ml were suspended into serum-free media. Mitochondrial membrane binding dye JC-1 (2.5 µg/ml, Abcam ab113850) was used for staining for 15 min at 37°C(Kim et al., 2018). Cells were washed with serum-free media after incubation and potential was measured on a flow cytometer with excitation of 488nm and emission at 530 and 590nm.

### ChIP Assay

Chip assay was performed essentially as described in this paper [34]. Briefly, worms of transgenic strain NL5901 were grown till desired developmental stage (L4). After harvesting the worms, they were crossed linked with 2% formaldehyde. Worms were sonicated to yield DNA fragments in the range of 200–500 bp. sonication cycle was adjusted to 20 sec on, 59 sec off at 55 percentage amplification for 5 minutes and 15 cycles. After sonication the DNA was pooled down with the help of respective antibodies. ChIP assay was performed in both estradiol untreated and estradiol treated groups using the H3K9ac (Abcam, ab16635) and H3K4me3 (Abcam, ab8580) Antibodies and rabbit IgG as negative control. This was followed by qPCR analysis of promoter region of *nhx-2*. IDT was used to design the primers for promoter amplification. Following primers were used to amplify the promoter region

*nhx-2* Promoter FP GCATGAGAGAGAGACGAGAGA

*nhx-2* Promoter RP AGGAAATCTGACACGCAAGAC

These promoter regions were further amplified with the help of qPCR of *the nhx-2* gene. The sequence of the promoter was obtained from Worm Base.

### Whole transcriptome analysis

To carry out this analysis, sample preparation for the homogeneous culture of NL5901, N2, VC363 worms were obtained by raising them on HT115 carrying the empty vector (control). Then this age synchronized L4 stage population was harvested for library preparation and proceeded for sequencing. The data was retrieved using the IlluminaHiSeq platform. The obtained reads were mapped onto the indexed reference genome of *Caenorhabditis elegans* (GCF_000002985.6_WBcel235) with the help of STAR v2 aligner [35] [36]. Following this, the differential expression analysis was conducted out using the DESeq2 package. Then this normalized test sample was analyzed w.r.t. the control sample. Here, the genes with absolute log2fold change ≥ 1.5 and p-value ≤ 0.05 were considered to be significant. The genes that conferred significant differential expression, were used for further analysis in the context of gene ontology (GO) and pathway enrichment analysis. For detecting the KEGG Pathway was performed using DAVID R package [37]. We also procured all the genes that are prominent in calorie restriction from the GenDR [38] database and plotted log2fold change of the differentially expressed genes in *nhx-2* knockout condition.

### Statistical analysis

The graphical data were presented as mean ± standard error of the mean. The data between the two groups were statistically analysed employing Student’s t-test by using GraphPad Prism v9.0 (GraphPad Software). p value < 0.05 was considered statistically significant for all comparisons.

## Results

### Preliminary RNAi screening of genes that delay reproductive senescence in context of various molecular events (alpha-synuclein aggregation, ROS levels, dopamine content) of neurodegeneration

Several genes are known to have a strong impact on reproductive health in *C. elegans*, for instance, RNAi of some of the genes delay reproductive senescence in *C. elegans* [18]. Taking these genes forward, we investigated the effect of their silencing in reference to various molecular events of neurodegeneration. Firstly, we assayed alpha-Synuclein expression, by employing transgenic strain of *C. elegans;* NL5901 [unc54p::alpha-Synuclein:: YFP + unc-119(+)], which expresses human alpha-Synuclein in muscles of the worm. We observed that control worms (NL5901) exhibited a mean fluorescence intensity of 19.31 ± 0.5957 arbitrary units whereas, RNAi of *nhx-2* exhibited the fluorescence intensity of 17.50 ± 0.5640 arbitrary units **(Figure.1a-b)**. Other genes also modulated the aggregation for instance *T04B2.1, VC27A7L.1, F25H8.1*, *C34D10.2* and *rskn-1* increased alpha-Synuclein expression and RNAi of genes *daf-2, C25G4.10, F20B10.3, F36F2.1* and *nhx-2* decreased alpha-Synuclein expression. *T04B2.1, VC27A7L.1, F25H8.1*, *C34D10.2* and *rskn-1* exhibited 22.03 ± 0.4810, 22.45 ± 0.5771, 24.45 ± 0.3211, 24.21 ± 0.8429, 22.23 ± 0.5497 and *daf-2, C25G4.10, F20B10.3, F36F2.1* and *nhx-2* exhibited a mean fluorescence intensity of 16.97 ± 0.4287, 16.85 ± 0.3447, 17.35 ± 0.4546, 16.11 ± 0.1126, and 17.50 ± 0.5640 arbitrary units respectively. **(Supplementary figure.1-2)**.

**Figure 1:**
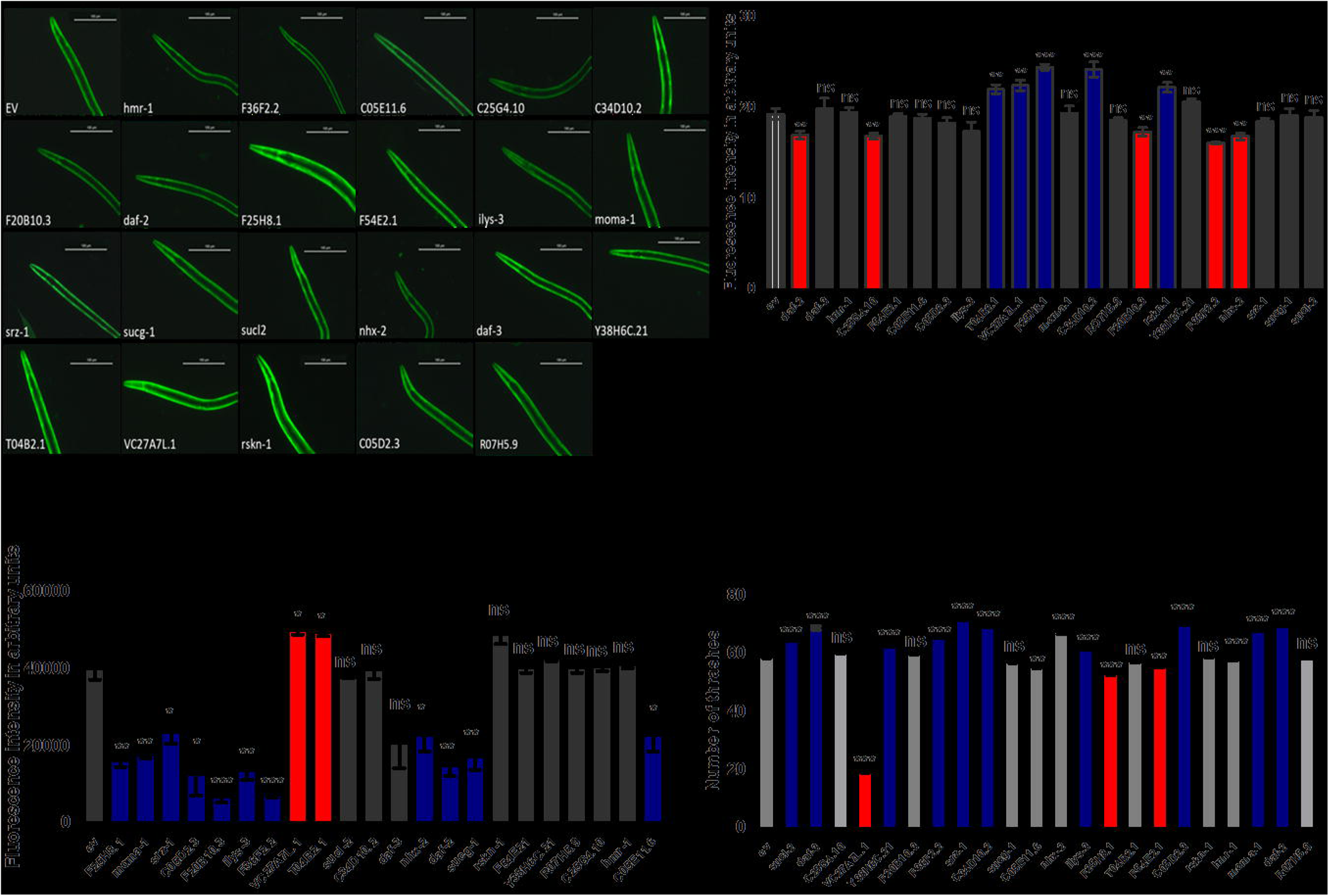
RNAi of *nhx-2* and its marked effect on various molecular events of neurodegeneration. (a) RNAi of *nhx-2* and its marked effect on □-Syn accumulation: The aggregation of □-Syn was assayed in NL5901 [unc54p::alpha-Synuclein:: YFP+unc-119(+)], a transgenic strain of *C. elegans* that expresses human □-Syn:: YFP transgene in their body wall muscles. All images used the same exposure time and gain settings; Scale bar=100µm. (b) Graphical representation for □-Syn aggregation in transgenic strain NL5901. (c) Effect of RNAi silencing of *nhx-2* on the relative formation of reactive oxygen (ROS) measured by H2DCFDA assay. H2DCFDA is a chemically reduced, non-fluorescent acetylated form of fluorescein which is readily converted to a green fluorescent form by the activity of ROS. An imbalance between the formation and transmission of ROS has been co-related with PD pathogenesis and can exacerbate its progression. (d) Effect of RNAi of *nhx-2* upon number of thrashes in transgenic strain NL5901 of *C. elegans*. The intensity of fluorescence was measured using ImageJ software and Significance was determined using Student□s t-test *p<0.05,**p<0.01,***p<0.001,ns-non-significant. (e)Marked effect of 6-OHDA on transgenic worm BZ555 in control EV and *nhx-2* knock down conditions respectively(A,B) A- in control worms there is evident degeneration of posterior dopaminergic neurons whereas in *nhx-2* RNAi condition this degeneration is rescued.

Age-associated neurodegenerative diseases are multifactorial. Many factors affect neuronal ailments including ROS, reactive oxygen species (ROS) alteration plays a very crucial role [39]. Hence, we studied the effect of RNAi of these genes on the level of ROS in the wild-type strain N2. We found that the level of ROS was 1.7 fold decreased in *nhx-2* RNAi condition **(Figure.1c)**, whereas, it was 2.4, 2.2, 1.6, 3.1, 5.9, 2.9, 5.2, 1.75, 2.6, 2.3 and 1.7 folds down-regulated in *F25H8.1, moma-1, srz-1, C05D2.1, F20B10.3, ilys-3, F36F2.2, nhx-2, daf-2, sucg-1, C05E11.6* respectively and 1.2 fold up-regulated in both *VC27A7L.1* and *T04B2.1* **(Supplementary figure. 3)**.

Aberrant motility is a hallmark feature of PD and is directly associated with the concentration of dopamine present in *C. elegans* [40]. To find out the effect of RNAi of these genes on the locomotion, we carried out a thrashing assay. Thrashing assay is useful for observation of the effect of loss of motor neurons. This assay was performed by counting the number of thrashes. We found that the number of thrashes were increased in RNAi group of *nhx-2* significantly. **(Figure.1d)** There were alterations of number of thrashes in other groups also for instance in *sucl-2, daf-3, C24G4.10, VC27A7L.1, F20B10.3, F36F2.2, moma-1,* and *CO5D2.3,* it was significantly increased whereas; in RNAi groups of *C34D10.2, T04B2.1, rskn-1* number of thrashes were decreased. **(Supplementary figure.4)**.

*While conducting our preliminary screening we observed that RNAi of nhx-2 significantly modulated all molecular events of neurodegeneration studied herein. We also found that nhx-2 knockdown not only delays RS but also increases longevity [41]. While studying estradiol treatment and associated effects we observed that nhx-2 is affected by estradiol treatment hence, it prompted us to study it further and we conducted further experiment to find the exact molecular mechanism for the same. The effects of nhx-2 knockdown on various molecular events have been given in the main image panel whereas; results of various molecular events, in RNAi condition of all 22 genes are given in respective supplementary figures 1-4*.

### RNAi of *nhx-2* induces Anti-alpha-Synuclein aggregation and neuroprotective effects

To verify our finding, we further performed an immunoblotting experiment using a fractionation method and employing an anti-aggregated alpha-Synuclein antibody. We found out that *nhx-2* decreases the aggregated form of alpha-Synuclein protein. *nhx-2* RNAi decreases aggregated alpha-Synuclein protein. The transgenic strain NL5901 fed with EV showed mean aggregation of 1.055 ± 0.05464 a.u. and worms fed with RNAi of *nhx-2* showed mean aggregation of 0.5072 ± 0.002500 a.u. which is significantly decreased. **(Figure.2a) (Figure.2b)**. We further checked its effects on neuronal health to investigate if it is eliciting any neuroprotective effect. To do that we procured another transgenic strain BZ555 that expressed GFP tagged with dopamine transporter and is expressed exclusively in dopaminergic neurons. We treated these worms with 6-OHDA that is known to degenerate dopaminergic neurons in both EV and *nhx-2* RNAi condition and we observed that RNAi of *nhx-2* is rescuing posterior dopaminergic neurons from degeneration **(Figure.1e)**

**Figure 2A.**
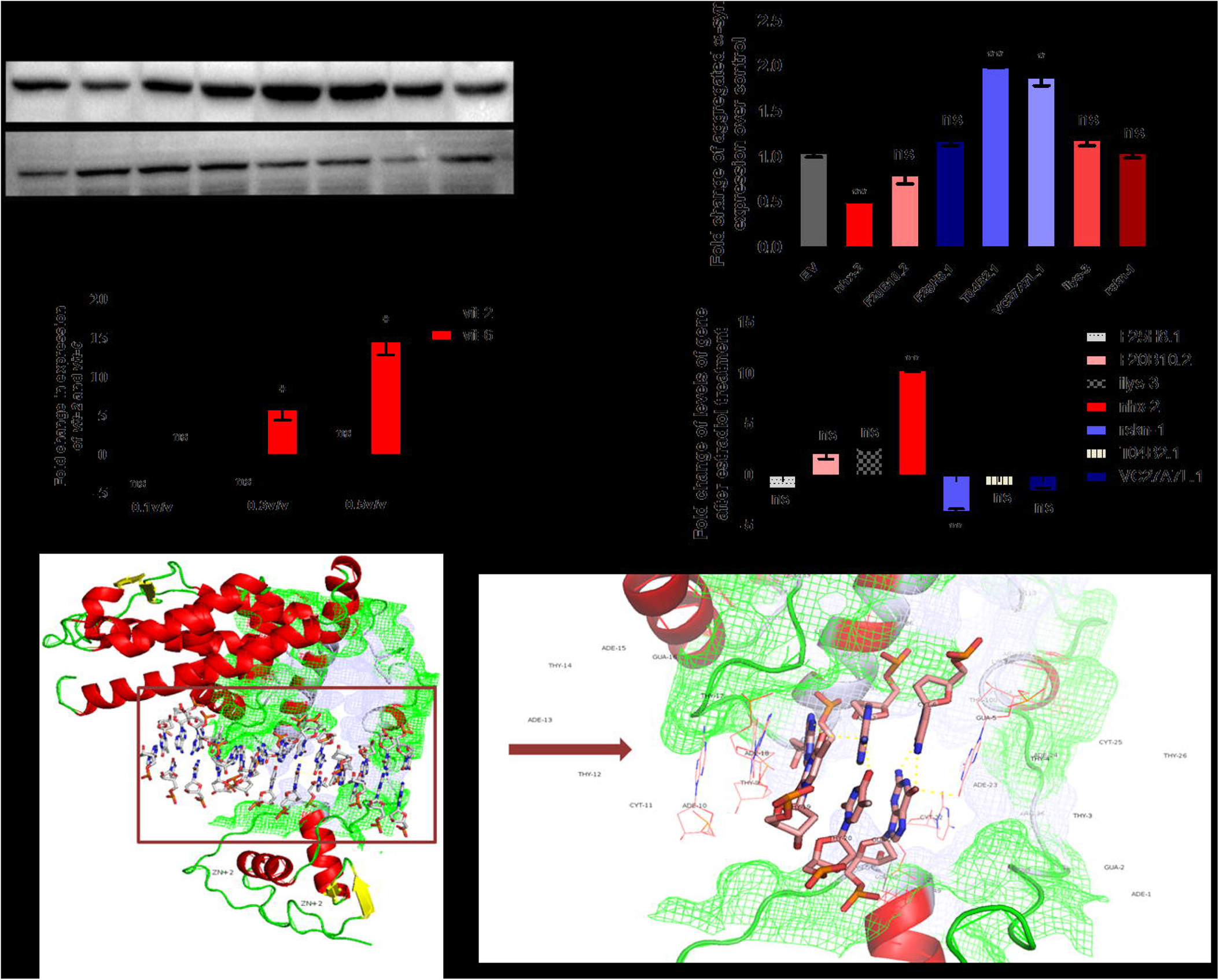
**Western blot and immunoblot analysis of alpha-Synuclein protein using anti aggregated alpha-Synuclein antibody:** (a-b): Represents the immunoblot analysis of aggregated alpha-Synuclein protein. Bands in first row represent soluble and aggregated alpha-Synuclein expression upon silencing of EV and *nhx-2* respectively, whereas bands in the second row represent control GAPDH, a housekeeping protein. Significance was determined using a Student□s t-test *p<0.05,**p<0.01,***p<0.001,ns-non-significant. (c-d): 17-β-estradiol treatment in *C. elegans* altered the transcripts of *nhx-2*. qPCR analysis of estradiol responding genes *vit-*2 and *vit-6* at different concentrations in wild type strain *p<0.05, ns-non significant. Real-time PCR analysis of screened genes after estradiol treatment over untreated control group in transgenic strain NL5901; Revealed estradiol treatment modulates *nhx-2*levels.Significance was determined using a Student□s t-test *p<0.05,**p<0.01,***p<0.001,ns-non-significant. (e) Interaction between Nuclear Hormonal Receptor and *nhx-2*, A) Docking of NHR-14 receptor-estradiol complex with *nhx-2* inverse repeat contig(AGTTGCAATACTA,1431-1443,5’-3’). B) There is an active interaction between the NHR-14 complex and DNA contig in DNA binding site. Participating residues are PHE,ARG,TRP,GLN,ASN,VAL,LEU,LYS,SER

### Estradiol receptor shows affinity, with the gene of *nhx-2* and Human 17-**β**-estradiol treatment alters mRNA levels of *nhx-2*

It is well known that estradiol plays a crucial role in the reproductive health but other functions of estradiol have also been discovered. In the previous finding, where reproductive senescence has been studied with respect to other health problems for instance osteoporosis, CVD and cancer; estradiol has demonstrated very critical role in all of them. Several studies show that estradiol plays a significant role in the maintenance and survival of various neurons [42]. The sexual dimorphism of predisposition of Parkinson’s disease in different genders and the timing of onset of reproductive senescence and neurodegeneration disease, presence of estradiol receptor in the brain in significant concentration and presence of estradiol receptor not only in nucleus but also in extra nuclear sites including synapse [43], all of these provide a clue that estradiol could play a key role in connecting both reproductive senescence and ND. Hence, we tried to check the effect of estradiol on the functioning of genes that is modulating various parameters of neurodegeneration.

#### a) *in silico* studies

It is well known that hormone receptors are cytoplasmic receptors. In the presence of their respective ligands, they move inside the nucleus and bind with the hormone response element and control transcription of various genes. While they function in a ligand-responsive manner, their binding domains look after varied functions. The functionality of hormone response elements (HRE) is highly dependent on the position of receptor and the consequent recruitment of the co-participants in the regulatory complex through which transcription is affected [44]. Translating the same insight to visualize the effect of *C. elegans* receptor of 17-β-estradiol NHR-14 and co integrator CBP-1 might affect the gene expression of *nhx-2*. To visualize if there is any binding affinity of the NHR-14 receptor with *nhx-2* gene’s inverse repeats; we performed docking studies. The results (**Table-2**) showed that this binding could be site-specific as it was showing affinity towards inverse repeat (AGTTGCAATACTA) from position 1431-1443 in gene *nhx-2*, the active residues in the binding pocket were PHE,ARG,TRP,GLN,ASN,VAL,LEU,LYS and SER respectively whereas, NHR-14 didn’t bind with another contig (GAAAAATTGTTCTA) from the position 1664-1667 of same gene. This obtained binding domain interaction was cross-validated using *her-1* which is known to have interaction with NHR-14, obtained from gene ontology in BioGrid [45]. The inverse repeat (TTTCATATCT) was taken from position 1578-1587 in gene. After docking the active residues participating in the binding pocket were; ARG,SER,VAL,TRP,GLN,LEU,GLU,TYR,THR,PRO, GLY, and ILE. It was observed that the binding sites i.e interacting pockets were located at DNA binding domain of NHR-14 for positive docked contigs of both the genes i.e *nhx-2* and *her-1*. Hence, it can be perceived from prior studies, literature and present docking that NHR-14 binds with the inverse repeats located upstream of the gene of interest which gives a clue that this gene could play a prominent and engaging role in Parkinson’s disease and its pathophysiology. **(Figure.2e)**.

### Docking summary of NHR-14-DNA contigs from genes *her-1* and *nhx-2* respectively: Table: 2

**Table 2:** Sequences of primers used in the real-time analysis and other experiments

| S. No. | Gene | Inverse repeat Sequence (5'-3') | Position | Receptor | Interacting residues number in the active site | Active residues | Group |
| --- | --- | --- | --- | --- | --- | --- | --- |
| 1 | *nhx-2* | AGTTGCAA<br>TACTA | 1431-1443 | NHR-14<br>estradiol<br>complex | 43,46,49,50,53,70,73,192,193,196,277,278,279 | PHE,ARG,TRP, GLN,ASN,VAL, LEU,LYS,SER | Hit Gene |
| 2 | *nhx-2* | GAAAAATT<br>GTTCTA | 1664-1667 |  | No interaction | No interaction | Negative control |
| 3 | *her-1* | TTTCATATC<br>T | 1578-1587 |  | 46,47,48,49,50,51,52,53,54,55,73,96,100,270,271,272,274,275,276,277,278,281,282,283,284,285,286,288 | ARG,SER, VAL,TRP, GLN,ASN, LEU, GLU, TYR, THR, PRO, GLY, ILE, | Positive Control |

#### *b)* Effect of human 17-β-estradiol on mRNA expression of *nhx-2*

To check whether human 17-β-estradiol affects the mRNA levels of *nhx-2* or not, we treated transgenic worms with 17-β-estradiol and checked the level of *nhx-2* along with other screened genes. To do so, we treated worms with different concentration of estradiol 0.1 v/v (volume by volume), 0.3 v/v and 0.5 v/v, and checked the level of estradiol-responsive vitellogenin gene and we found out that concentration 0.5 v/v increased the expression of both *vit-2* and *vit-6* genes, any further increase in concentration decreased *vit-2* expression **(Figure.2c)**. We checked the transcription levels of all screened genes that altered molecular events to neurodegeneration to be sure that estradiol is working in the worms. We found that upon 17-β estradiol treatment, there were two genes whose mRNA levels were modulated namely; *nhx-2* and *rskn-1*. So, it validated our *in-silico* data. The level of *nhx-2* was up-regulated whereas; the level of *rskn-1* was down-regulated **(Figure.2d)**. Since RNAi of *nhx-2* was the one that modulated all the parameters of ND’s and *rskn-1* did not show any such effects, we hypothesized that reduced level of *nhx-2* could play a crucial role in the maintenance of neuronal health. Hence, it prompted us to extent our further work to decipher the exact mechanical pathway involved in the associated effects.

### ChIP analysis ascertains that estradiol treatment induces H3K4me3 and H3K9 acetylation activation of the promoter of endogenous *nhx-2 gene* thereby increasing its expression level

Once we deciphered that estradiol increases the transcription levels of *nhx-2*, we performed an *in silico* analysis and found that estradiol receptor NHR-14 shows significant affinity with the *nhx-2* gene. To assess that estradiol treatment leads to *nhx-2* induction by directly modifying the chromatin at its promoter, we analyzed the histone acetylation level at *nhx-2* promoter through chromatin immunoprecipitation (ChIP) assay, hence we investigated whether estradiol treatment is altering the epigenetic state of *nhx-2* promoter region. Prior to our study at the specific promoter, we first investigated the effect of estradiol treatment on histone acetylation at the global genomic level, while the global expression of H3K9Ac was found to be significantly increased, we found no significant alteration of H3K4me3 **(Figure.3b)**. We further checked the occupancy of H3K4me3 and H3K9 acetylation activation marker of *nhx-2* promoter region after estradiol treatment with the help of ChIP analysis. The chromatin was first probed with specific antibodies followed by validation with specific primers. Our results demonstrated that human estradiol treatment gives rise to enhanced occupancy of both H3K4me3 and H3K3 acetylation markers on the promoter region of *nhx-2* thereafter; resulting in their increased expression. H3K9 acetylation marker was 2.167 fold and H3K9me3 marker was 3.8 fold upregulated on the promoter region of *nhx-2* upon estradiol treatment. Where untreated (UT) condition showed a mean fold change of 0.5374 ± 0.002896 and1.586 ± 0.06830, the estradiol treated worms showed amean fold change values of 1.164 ± 0.04872 and 5.864 ± 0.07550 of H3K9 ac and H3K4me3 activations respectively. Hence, our data provides evidence that estradiol treatment increases the expression of sodium-proton pump in gut via inducing epigenetic modification on the promoter region of the *nhx-2* gene **(Figure.3e)**.

**Figure 3:**
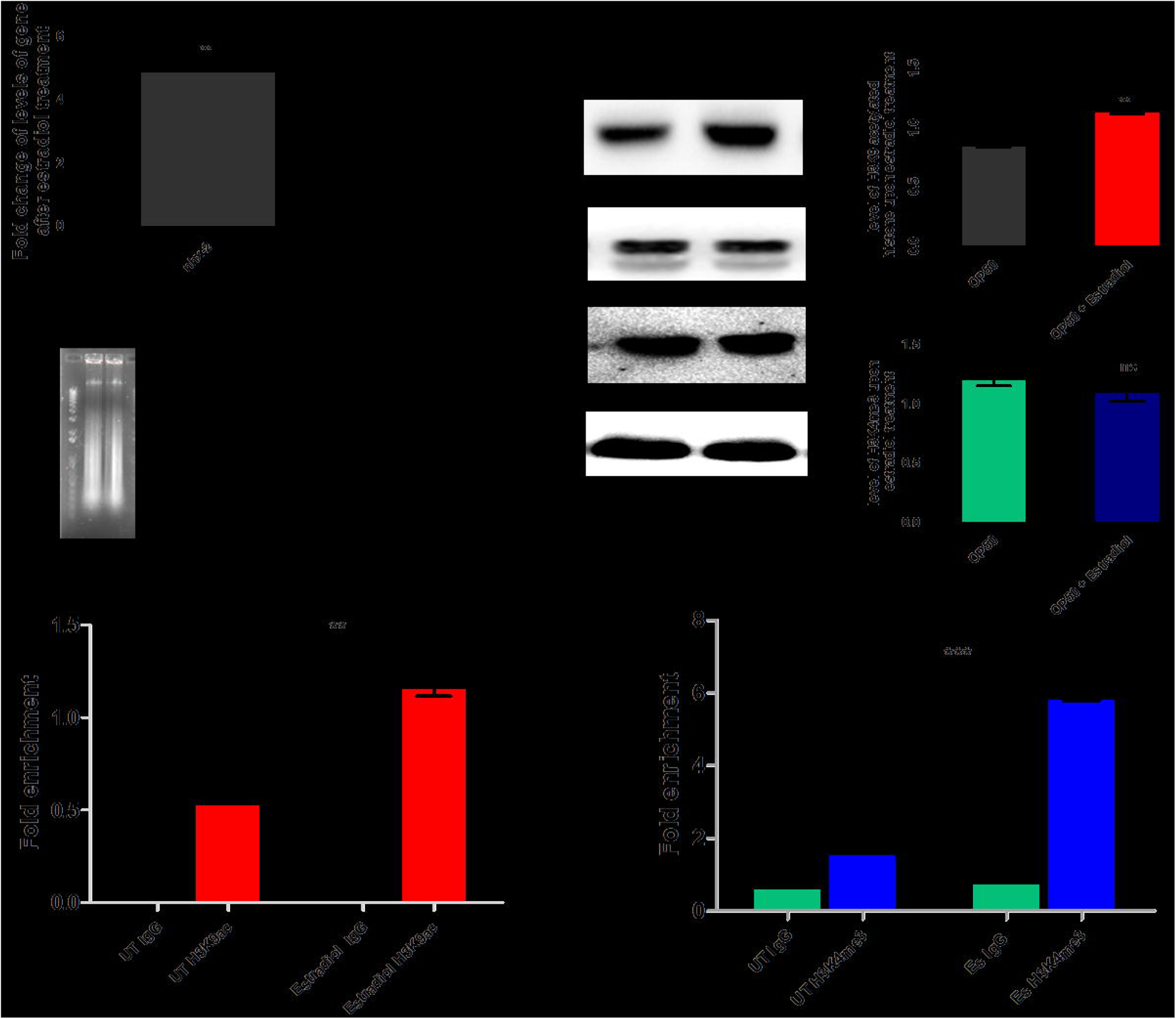
ChIP analysis demonstrating estradiol treatment increases the occupancy of H3K9ac and H3K4me3 activation marker on the promoter region of *nhx-2*. (a) qPCR analysis demonstrating significant upregulation of *nhx-2* upon estradiol treatment, data is presented as ± SEM (n=2), Significance **p<0.01 (b) This image shows the effect of estradiol treatment on H3K9 acetylation at the global genomic level. While the global expression of H3K9ac was found to be significantly increased we found no significant alteration of H3K4me3. The significance level was determined using ImageJ analysis followed by graph pad where significance levels were calculated as **p<0.01, ns-non-significant (c) The image represents different sizes of DNA fragments (200bp) after sonication with the help of agarose gel electrophoresis. (d-e) Occupancy of H3K9ac and H3K4me3 activation markers at the promoter region of the *nhx-2* gene were evaluated with the help of ChIP analysis, using the H3K9Ac and H3K4me3 antibodies in both untreated and estradiol treated groups followed by qPCR analysis using specific primers of the promoter regions. Data are presented as ± SEM (n=2), Significance **p<0.01,ns-non-significant.

### *nhx-2* upregulation does not amend alpha Synuclein aggregation any further

With the help of immunoblot experiment we checked the level of aggregated alpha Synuclein in both downregulated and upregulated (induced via estradiol treatment) conditions of *nhx-2* and we found that although RNAi of *nhx-2* decreases the alpha-Synuclein aggregation significantly (where control NL5901 worms showed aggregation of 1.050 ± 0.05000 arbitrary units, RNAi of *nhx-2* showed aggregation of 0.5547 ± 0.05000 arbitrary units), increased expression of *nhx-2* via estradiol treatment, does not alter the aggregation any further.**(Figure.4Aa)**

**Figure 4A:**
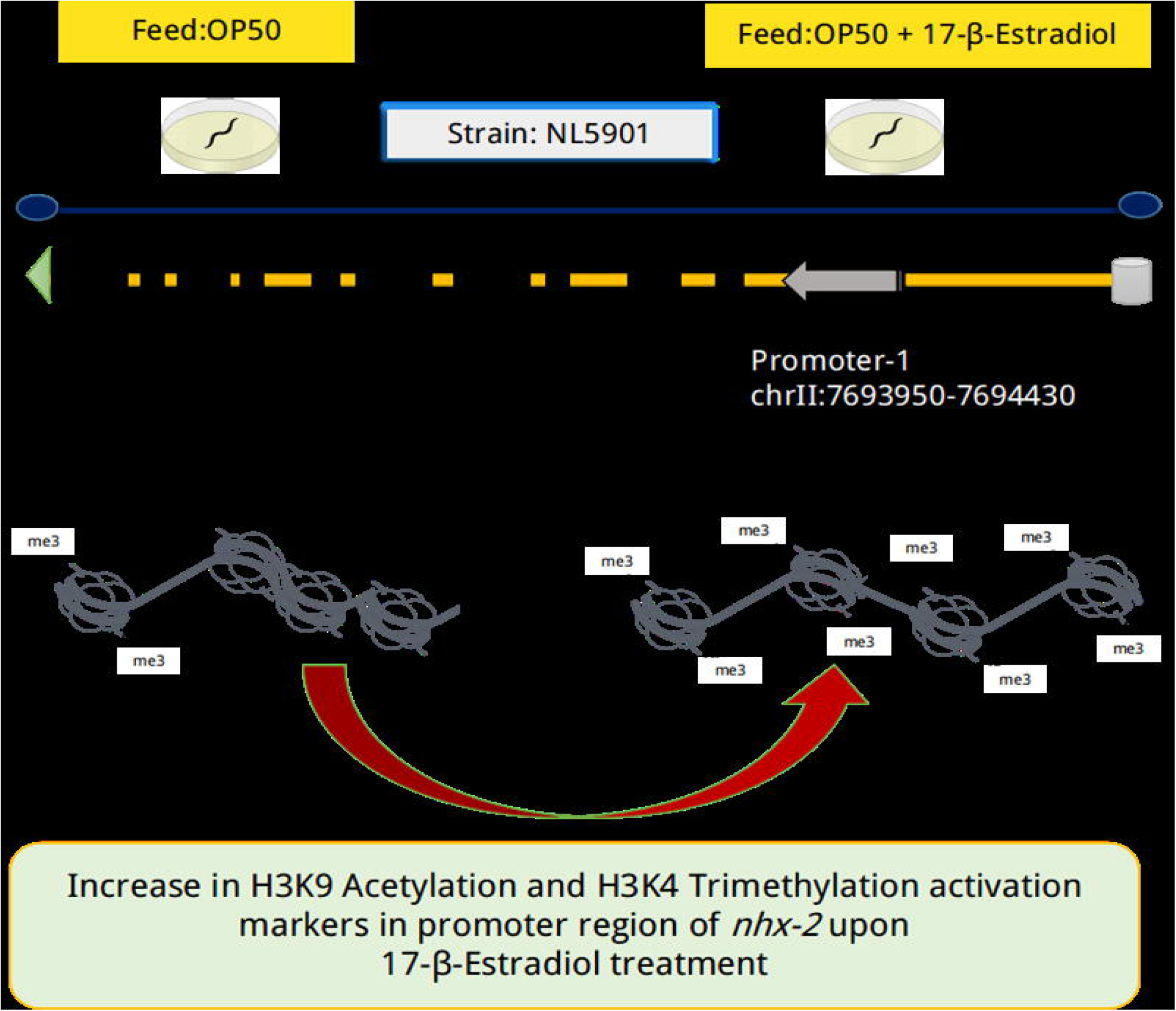
**Estradiol treatment increases *nhx-2* expression although it doesn’t give rise to increased alpha-Syn aggregated protein and RNAi/Knockout of *nhx-2* alters food absorption, accelerates *sir-2.1* expression and functionality.** (a)This figure represents results of the immunoblot experiment in which bands of the first row represent alpha-Syn aggregated protein in control, *nhx-2*-RNAi and worms treated with estradiol respectively whereas bands in the second row represent housekeeping protein GAPDH. Significance was determined using a Student□s t-test *p<0.05, ns-non-significant. (b) This image panel represents fluorescent OP50 in the gut of NL5901 and Knockout strain of *nhx-2*: VC363.A-B in the panel represents anterior and posterior parts of the worm NL5901 and C-D in the panel represents anterior and posterior parts of the KO strain of *nhx-2* VC363. Arrows in image C-D points unabsorbed food that remained in the gut, thereafter the entire gut is showing fluorescence. (c) This figure represents qPCR analysis of mRNA levels of *sir-2.1* upon *nhx-2* RNAi. Significance was determined using a Student’s t-test **p<0.001. (d) RNAi of *nhx-2* increased the reproductive life span by five days in transgenic strain NL5901 but this alteration of the reproductive life span was not shown in the RNAi condition of sir-2.1.

### RNAi of *nhx-2* decreases the absorption of food from the gut and mimics calorie restriction thereby; exerting its effect on SIR-2.1 expression

In order to find out exact pathway of how RNAi of *nhx-2* affects various parameters of NDs, we mined the available literature on the subject, and we found that knockdown of *nhx-2,* which is expressed in the apical membrane of epithelial cells, not only alters the intestinal pH, it is also associated with *opt-2* which, is an oligopeptide transporter that helps in food absorption [41]. Hence, we hypothesized that alteration in *nhx-2* expression might interfere with the food absorption and to ratify our hypothesis, we carried out food absorption studies employing an *E. coli* strain; OP50 tagged with green fluorescent protein (GFP). These fluorescent bacteria were fed to NL5901 and *nhx-2* knockout worms (VC363) towards assaying the level of food absorbed within them. We found that in *nhx-2* knock out condition significant amount of food remained in gut and rectum region in comparison to NL5901. Hence, less food absorption exerts an effect like calorie restriction (CR), and probably this is how its RNAi is acting as a neuroprotectant. To test our hypothesis, we carried out further experiments. **(Figure.4Ab)**.

### Whole transcriptome analysis, corroborates the CR effects elicited by silencing of *nhx-2*

In order to substantiate and further validate our previous finding that silencing of *nhx-2,* acts as CR mimetic, we performed whole transcriptome analysis in knock out strain of *nhx-2*; VC363 and checked the transcriptional state of various marker genes of CR. Our data demonstrated a comprehensive information about the functions associated with *nhx-2* and its role in various biological processes. The gene ontology obtained from the R package has shown that knock out of *nhx-2* alters biological processes like oxidation-reduction process, ion transport, cell division, innate immunity response, steroid mediated signaling pathway, gonad development, protein transport, cell development, check point regulation in cell cycle, gonad development and metabolic processes.We observed that genes that are being deferentially expressed are component of plasma membrane, cytoplasm, and nucleus. We further checked the differential gene expression of marker genes of CR. As described here in GenDR database, there are 62 marker genes of CR and our data revealed differential expression of 61 hallmark genes out of 62, in knock out condition of *nhx-2*. **(Figure.4B)**

**Figure 4B:**
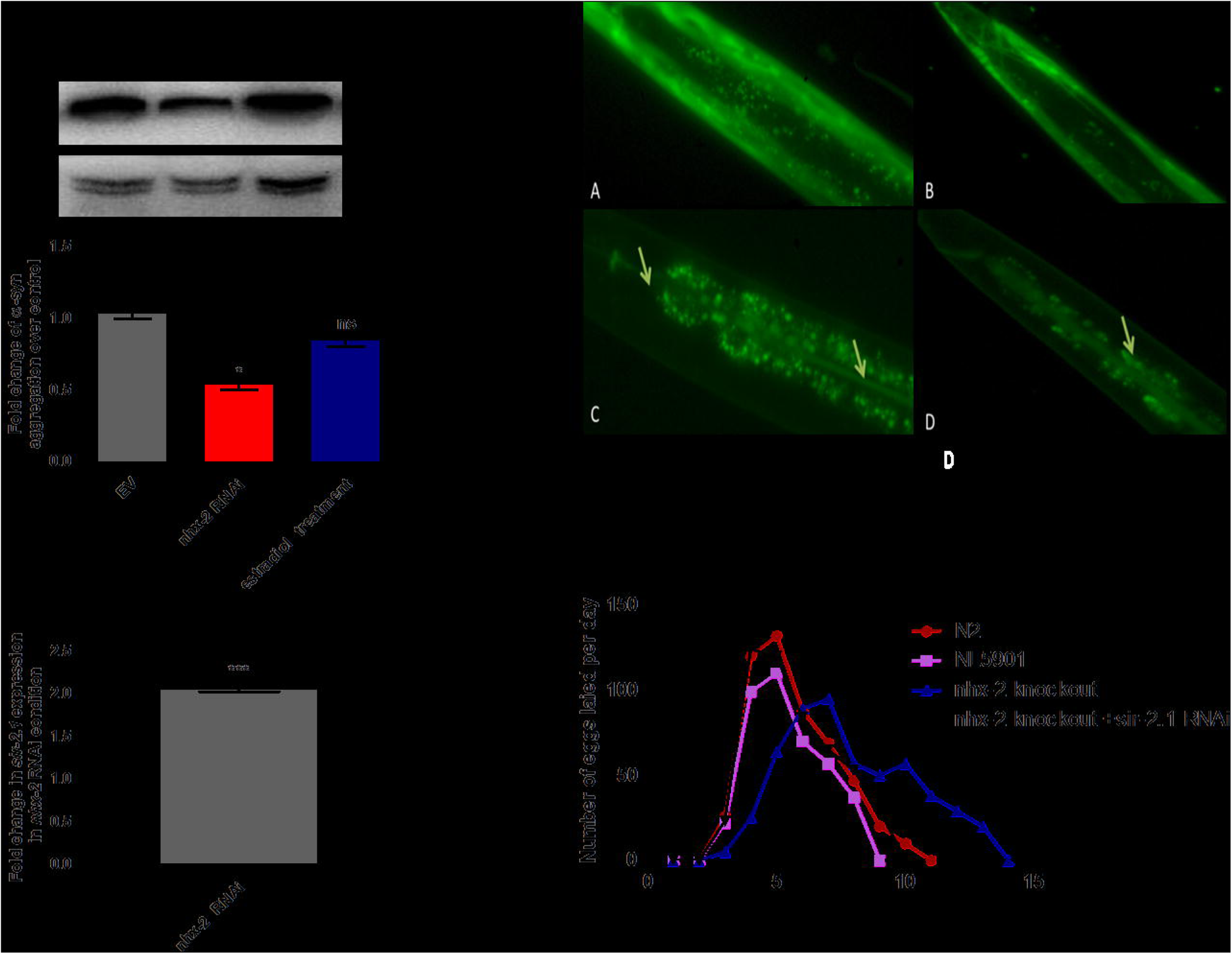
**Whole transcriptome analysis corroborates that silencing of *nhx-2* acts as genetic mimetic of calorie restriction.** (a) Differential expression of the genes involved in dietary restriction was obtained from GenDR, herewith representative graph displays significantly upregulated and downregualated genes in *nhx-2* knockout condition; in which *eat-2* was significantly downregulated. This illustrated the underlying reason for calorie restriction mimic. (b) HeatMap representation of significant genes from whole transcriptome data. (c) Volcano plot of differential expression genes, where red represents upregulated, green represents downregulated and grey represent neutral genes.

After finding that *nhx-2* knockout/RNAi mimics CR and that it also exerts neuroprotective effect, we speculated that it might be affecting SIR-2.1 levels because based on previous findings, it is been suggested that CR exerts its neuroprotective effect in presence of SIR-2.1. Hence, we checked the mRNA levels of *sir-2.1* in RNAi conditions of *nhx-2* and we found that in *nhx-2* silenced worms the levels of SIR-2.1 were more than four fold upregulated **(Figure.4Ac)**. We now know that *nhx-2* RNAi gives rise to increased mRNA levels of *sir-2.1* but we also checked, whether this gets translated into increased protein levels of SIR-2.1 thereafter; alters the function of SIR-2.1. In order to investigate this, we performed genetic crossing in order to estimate protein expression as well as fecundity assay to access the coherent networking of *nhx-2* with *sir-2.1*.

### RNAi of *nhx-2* extends reproductive life span only in presence of SIR-2.1

In order to ascertain the regulatory networking of NHX-2 and SIR-2.1, we employed fecundity assay in various treatment conditions. This assay demonstrated reproductive window of worms in various groups studied herein; for instance, N2 as control group, NL5901 as pathological strain, knock out strain of *nhx-2*; VC363 and one more group wherein; in knock out strain of *nhx-2*, we performed RNAi induced silencing of *sir-2.1*. This assay not only demonstrated reproductive window of worms but also reflected the brood size of worms. We observed that wild type worms lay eggs till day 6, worms of NL5901 stain demonstrated egg laying till five days whereas; worms of *nhx-2* knockout strain, lay eggs till day 10 hence, demonstrated a significant delay in RS, precisely four days from control. However, when *sir-2*.1 is silenced in the knock out strain of *nhx-2,* worms demonstrated egg laying till seven days. The data herein; demonstrated that the delay in reproductive health was nonsignificant, when *sir-2.1* was silenced in background of *nhx-2* knockout. Although there is a huge shift in reproductive window of *nhx-2* knock out worms, we observed that there was no alteration in brood size of worms of the all group. This data strongly argues that the effects induced by *nhx-2* silencing are driven by *sir-2.1* **(Figure. 4Ad).** The data presented herein; corroborates the prominent role of NHX-2 in the entire network involved in calorie restriction and it’s the underlying mechanism that in return is effecting on worm’s health.

### Creation of a transgenic strain that expresses ubiquitous mCherry SIR-2.1 and alpha-Synuclein intact with its muscles signifies that *nhx-2* RNAi alters the expression of SIR-2.1

To substantiate our findings that knockdown of *nhx-2* exerts its neuroprotective effect via up-regulating the expression of SIR-2.1, we created a transgenic strain that expresses SIR-2.1 ubiquitously and alpha-Synuclein intact with its muscles. We performed several genetic crosses along with several backcrosses between transgenic strain UL2992 [sir-2.1::mCherry + rol-6(su1006)] that expresses mCherry tagged SIR-2.1 and transgenic strain NL5901 that expresses alpha-Synuclein intact with its muscles. With the help of genetic and backcross, we have constructed a strain that expresses mCherry tagged SIR-2.1 ubiquitously and YFP alpha-Synuclein in the muscles. We further performed *nhx-2* RNAi in this strain and we observed that knockdown of *nhx-2* in these worms, decreases alpha-Synuclein expression and increases SIR-2.1 expression. This experiment hence further substantiated our previous finding that *nhx-2* RNAi exerts its neuroprotective effect via modulating the expression of SIR-2.1 **(Figure.5)**.

**Figure 5:**
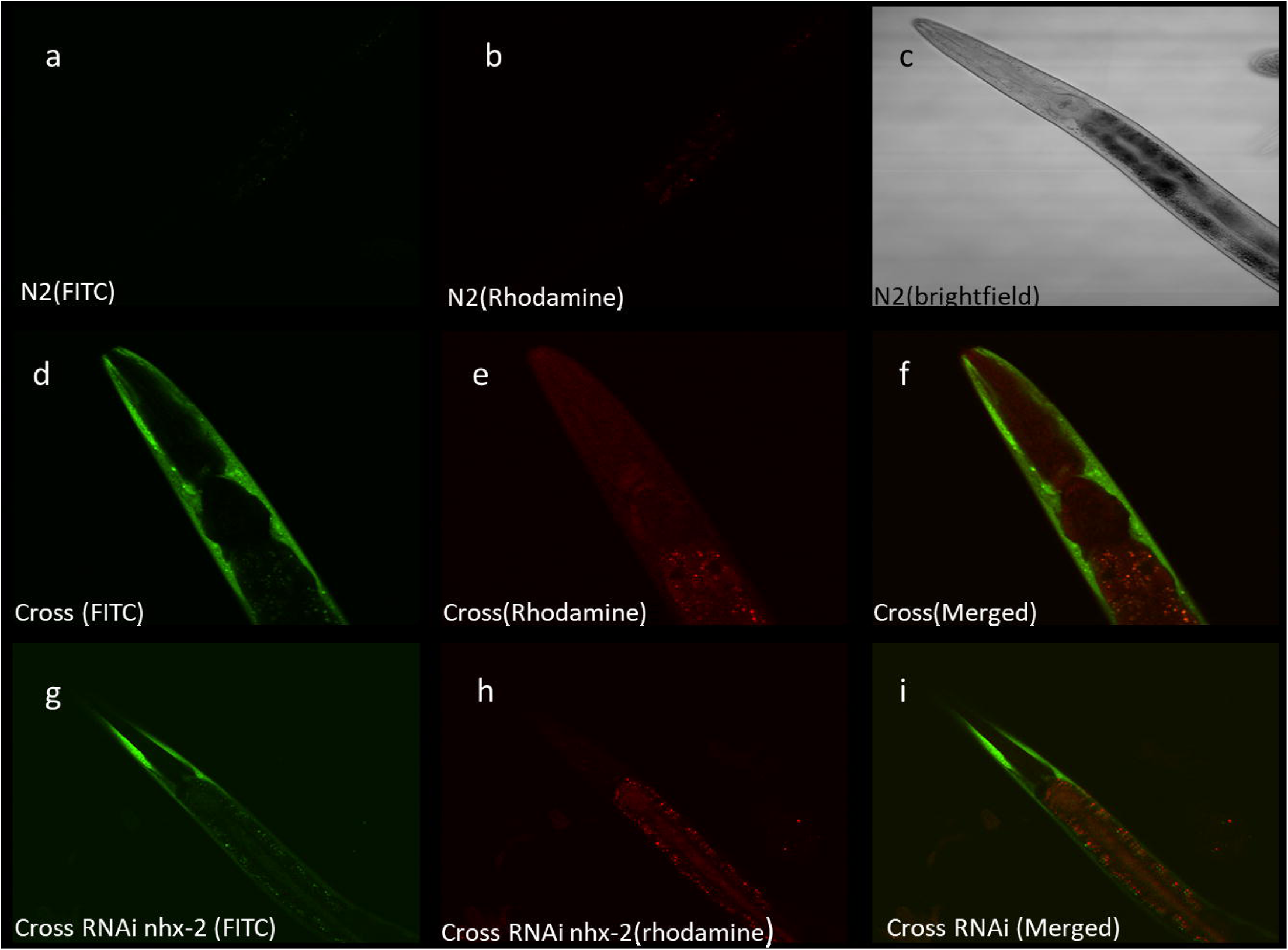
Creation of transgenic strains that express SIR-2.1 ubiquitously and alpha-synuclein in its muscles revealed that *nhx-2* RNAi decreases alpha-synuclein via up-regulating the expression of SIR-2.1. (A-C) Confocal images of the wild type strain in FITC, Rhodamine and brightfield (D-F) Confocal images of crossed EV worms in FITC, Rhodamine and merged image (G-I) Confocal images of crossed worms given *nhx-2* RNAi treatment in FITC, Rhodamine and merged image

### RNAi of *nhx-2* improves mitochondrial functioning and restores mitochondrial membrane potential in PD model of *C. elegans*

In order to perform pathway analysis and to decrypt the exact molecular processes that are eventually getting attuned by silencing of *nhx-2 w*e performed qPCR analysis taking those gene into consideration that are admitted markers of various molecular pathways, for example; *lin-45, jkk-1* are well recognized apoptotic markers *lgg-1, atg-5, vps-34 are* autophagy markers, whereas*; atp-2, gas-1*, and *cco-2* are genetic markers of mitochondrial functioning. Our data suggested that mRNA levels of genes of mitochondrial functioning were being modulated. We found that *atp-2* which encodes the beta subunit of the soluble, catalytic F1 portion of ATP synthase (mitochondrial respiratory chain [MRC] complex V) was 17.36 fold up-regulated whereas; *gas-1* whose expression is controlled by *atp-2* via feedback inhibition [46] also down-regulated by 2.06 folds **(Figure.6a)**. The above data sets a clue, indicating that if knockdown of *nhx-2* is modulating a component of ATP synthase; it might also be affecting the overall production of ATP and the number of mitochondria. To check their levels, we performed ATP assay and mitotracker imaging to know the content of healthy mitochondria in the knockdown condition of *nhx-2*. We found that the ATP level was 1.33 fold up-regulated **(Figure.6b)** and several mitochondria appeared to be increased significantly in *nhx-2* knockdown condition **(Figure.6c)**. Now the question was how does knockdown of a proton pump can modulate ATP levels and mitochondrial content? To know the answer, we checked the change in potential of the mitochondrial membrane which is responsible for proton gradient and thereby ATP production as well. To check the modulation in mitochondrial membrane potential of each cell we first of all isolated cells of *C. elegans* following the method described earlier. We checked out the change in mitochondrial potential in control wild-type worms, in PD model NL5901 and worms treated with knockdown of *nhx-2* in PD model NL5901 with the help of flow cytometry and we found out that RNAi of *nhx-2* is driving the membrane potential towards wild-type N2 strain by restoring mitochondrial depolarization **(Figure.6d) (Figure.6e)**.

**Figure 6:**
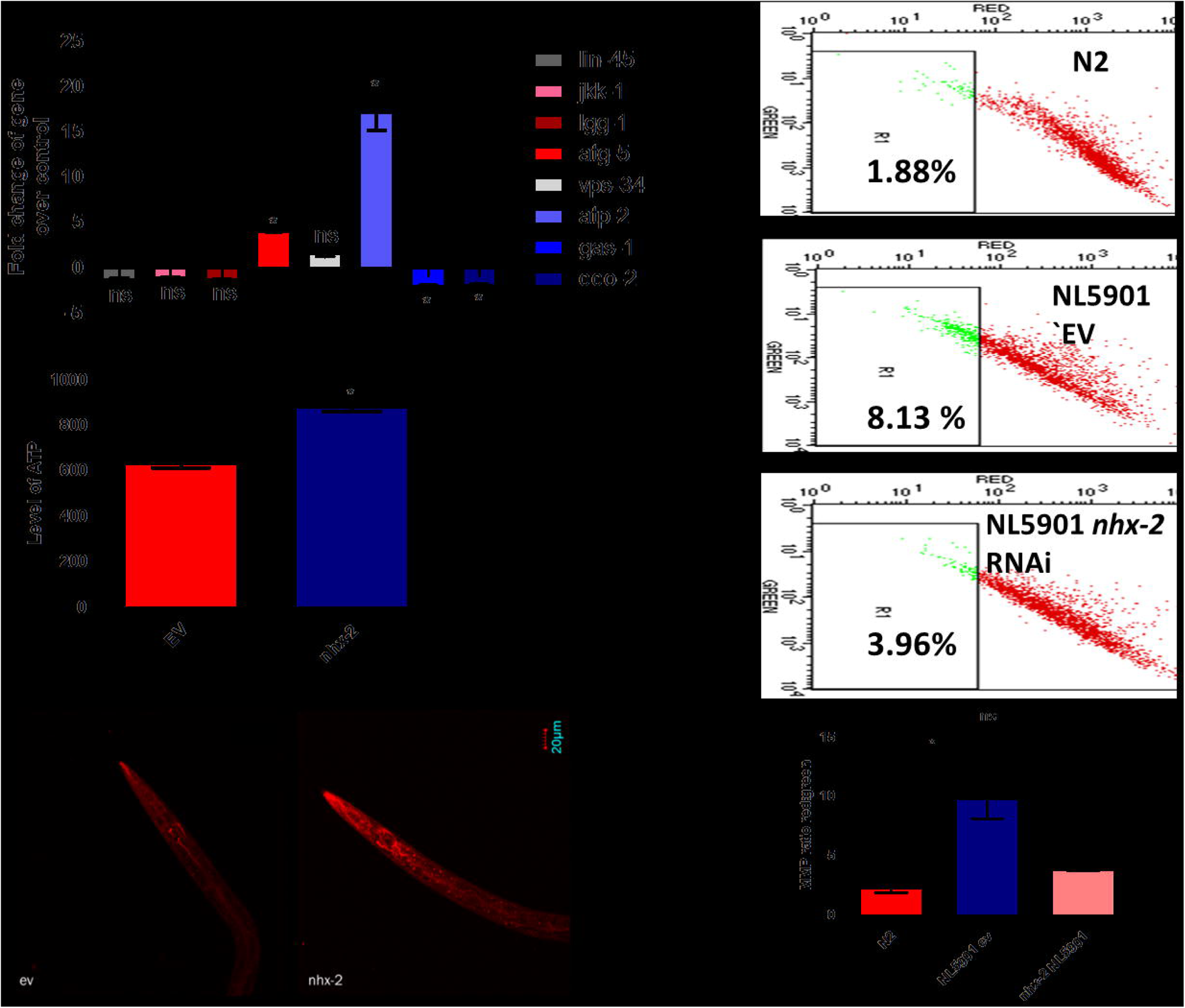
*nhx-2* RNAi improves overall mitochondrial health and restores the mitochondrial membrane depolarization in transgenic strain NL5901. (a) qPCR analysis of selected genes of various pathways in transgenic model NL5901. The level of gene *atp-2* is significantly up-regulated whereas the level of genes gas-2 and cco-2 is significantly down-regulated. *p<0.05, ns-nonsignificant) ATP measurements: The level of ATP was significantly up-regulated in worms with *nhx-2* RNAi treatment. Significance was determined using a Student’s t-test (*denotes p<0.05 as compared to the wild-type,***denotes p<0.001 as compared to the wild type). Data represent mean±SEM. c) The effect of knockdown of *nhx-2* on mitochondrial content in transgenic strain NL5901 of *C. elegans* that expresses human □-Syn:: YFP transgene in their body wall muscles. All images used the same exposure time and gain setting, Scale bar=20µm.d) RNAi of nhx-2 restores depolarization of mitochondrial membrane; 1) Mitochondrial membrane potential [MMP] of wild type N2 strain, in wild type N2 strain the j-aggregates are lower which shows that the membrane is depolarised, 2) MMP of transgenic worms NL5901 expressing □-Syn. The mitochondrial membrane is polarized which is indicated by higher j aggregates.3) RNAi of *nhx-2* restores the depolarization of mitochondrial potential in transgenic strain NL5901 as evident from the image it decreases j aggregates significantly. e) MMP ratio of red vs green cells to measure the MMP change in wild type, EV, NL5901, and *nhx-2* RNAi conditions. Significance *p<0.05,**p<0.001.

These experiments suggested that RNAi of *nhx-2* is exerting its neuroprotective effect by mimicking calorie restriction and improving mitochondrial membrane potential.

## Discussion

Obtaining an in-depth understanding of processes related to physical, cellular and molecular ageing has become imperative, as global life expectancy witnesses a significant extension. A critical dimension that is associated with ageing is; “decline in reproductive capacity” or “reproductive senescence (RS)” When reproductive senescence sets in, it gives rise to several changes that affect the susceptibility of aged individuals towards several health conditions, for instance, cardiovascular diseases, osteoporosis, and neurodegenerative diseases. One major change that immensely concerns neuronal health is; the change in blood levels of estradiol. The presence of estradiol receptor GPERs in brain cells, sexual dimorphism in the prevalence of different neurodegenerative diseases and other facts such as observational studies indicating estrogen use in postmenopausal women and its association with reduced (25–70%) risk of neurodegeneration, prove that estradiol is an important factor in neuronal health and integrity [47].

In the present study, we have explored the role of such genes that are known to extend reproductive span in the context of neurodegeneration. There are several genes in the genome of *C. elegans* which, upon silencing delay their reproductive senescence. This particular study has been conducted employing the Ahringer *C. elegans* RNAi feeding library, wherein; we have performed RNAi induced silencing of 22 genes, whose RNAi clones are available in the Ahringer library and studied associated effects in context of neurodegeneration. While performing our preliminary screening, we studied these genes with respect to various molecular events of neurological ailments such as alpha-Synuclein aggregation, ROS production, and dopamine levels through aversion assay. We further investigated whether human estradiol induces any modulation in the transcription levels of the screened genes to find out their relevance in neurodegenerative disease vis-à-vis reproductive aging. While performing our systematic screening we kept two parameters in consideration 1) genes that consistently and significantly modulate the molecular events associated with neurodegeneration studied herein; including protein aggregation, ROS levels, mobility and 2) genes that are being modulated by the presence/absence of human estradiol. These studies led us to identification of *nhx-2*, a proton sodium anti-porter, which is expressed exclusively on the apical plasma membrane of the intestine. *nhx-2* demonstrated strong consistent results modulating all parameters studied herein, and it is transcriptionally modulated by human estradiol treatment. With the help of in silico analysis, we observed that NHR-14 which is the receptor of human estradiol in *C. elegans* shows affinity with the invert repeats present in the promoter region of the *nhx-2* gene. NHR-14 is a receptor that is present in the cytoplasm in the cell but when it binds with its ligand, it gets translocated into the nucleus and modulates the transcription of several genes. NHR-14 is a receptor of human estradiol and we found that it demonstrates the affinity to bind with the invert repeats of promoter region of the *nhx-2* gene. In order to ascertain our findings, we performed ChIP analysis to check the epigenetic modification on the promoter region of *nhx-2* gene induced by estradiol treatment. We checked the occupancy of H3K4me3 and H3K9 acetylation activation marker designing specific primers of the promoter region of *nhx-2* gene after estradiol treatment. We found that estradiol treatment gives rise to increased occupancy of both H3K4me3 and H3K9 acetylation marker in the promoter region of the *nhx-2* gene. Our data strongly argues with our previous finding and signifies that estradiol governs the levels of this proton pump in gut.

Our knowledge of the biological role of sodium-proton transporters in the intestine has been around pH maintenance, food absorption, lipid storage and larval development [41]. In the present study, we observed the marked knockdown effect of the sodium-proton antiporter on neurodegenerative disease in *C. elegans*. Our observations substantiate previous findings that the loss of *nhx-2* gives rise to alteration of pH which leads to a phenotype that mimics starvation and we further found that the effect exerted via RNAi of *nhx-2* is governed by SIR-2.1. We observed that an increased healthy life span due to calorie restriction (CR) elicits strong neuroprotective effects. Having observed that RNAi of *nhx-2* is mimicking calorie restriction, we tried to find out the exact molecular mechanism involved in neuroprotection. To do that we performed whole transcriptome analysis in knock out strain of *nhx2*; VC363. This study gave us a comprehensive information regarding the biological role of *nhx-2*. To decipher the effects associated with *nhx-2* and its networking with CR induced gene modulation, we checked the transcriptional state of genes that are enlisted as recognized markers of CR in worms at GenDR. GenDR had around sixty-two genes that are associated with the dietary restriction, out of these sixty-two genes sixty-one were found to be differentially expressed in the *nhx-2* knockout condition, suggesting the prominent role of NHX-2 in the entire network involved in calorie restriction and it’s the underlying mechanism that in return is affecting neuronal health. In one of our previous studies, we have found that calorie restriction and neuroprotection are linked together via SIR-2.1[20]. Silent information regulator 2 (Sir-2.1) proteins, or sirtuins, are protein deacetylases that are found in various organisms, from bacteria to humans. In our previous work, calorie restriction was found to show its protective effect only in the presence of *sir-2.1*. Keeping that in mind we hypothesized that RNAi of *nhx-2* must be showing its effects via *sir-2*.*1*. To test this, we have investigated the interplay between *nhx-2* and *sir-2.1* using a combination of RNAi, real-time PCR studies, western blot, pathway mapping via bioinformatics tool GENEMANIA and genetic crossing. First of all, we checked whether there is any alteration in the *sir-2.1* levels in the knockdown condition of *nhx-2*. To answer this query, we performed real-time PCR analysis to quantify the levels of *sir-2.1* in *nhx-2* RNAi condition and we observed that in *nhx-2* RNAi condition, the level of sir-2.1 was upregulated significantly. In order to check whether this transcriptional elevation gives rise to increased protein levels of SIR-2.1 we performed a genetic cross between two transgenic strains UL2992 [sir-2.1::mCherry + rol-6(su1006)] that expresses mCherry tagged SIR-2.1 ubiquitously and transgenic strain NL5901 that expresses alpha-Synuclein intact with its muscles. With the help of a genetic crossing followed by the back cross, we have constructed a strain that expresses mCherry SIR-2.1 ubiquitously and possesses YFP alpha-Synuclein expression in the muscles. We further performed *nhx-2* RNAi in this strain and we observed that knockdown of *nhx-2* in these worms decreases alpha-Synuclein expression and increases SIR-2.1 expression. This experiment hence further substantiated our previous finding that *nhx-2* RNAi exerts its neuroprotective effect via increasing the expression of SIR-2.1. It was clear from our previous experiments that sir-2.1 mRNA levels are up-regulated in *nhx-2* knockdown condition but whether RNAi/knockout of *nhx-2* is affecting the function of SIR-2.1 was yet not clear. To investigate this, we procured knock out strain of *nhx*-2 and we performed fecundity assay in RNAi background of *sir-2.1* and we found that reproductive health extension induced by RNAi of *nhx-2* is SIR-2.1 driven.

To decipher the specific mechanistic pathway, we performed pathway analysis with the help of real-time PCR analysis and checked the mRNA levels of various genes critical for several cellular regulatory processes. We found out that amongst all, the genes of mitochondria for example : *atp-2, cco-2,* and *gas-2* were significantly modulated. It is also known that calorie restriction via SIRT-2.1 alters mitochondrial biogenesis with increased respiration and enhanced ATP production (Nisoli et al., 2005). *nhx-2*, being a proton pump and altering the levels of *sir-2.1* might as well affect overall mitochondrial health. Hypothesizing that we checked various parameters of mitochondrial health in *C. elegans* including the amount of ATP, number of mitochondria with the help of mitotracker and we have for the first time in *C. elegans* checked the mitochondrial membrane potential of individual cells of *C. elegans* in the knockdown condition of *nhx-2*. We found out that knockdown of *nhx-2* not only elevated the number of mitochondria consequently levels of ATP but also it restored the depolarization of mitochondrial membrane potential in transgenic strain NL5901.

Our observations led us to the conclusion that estradiol connects neuronal health with gut homeostasis via regulating *nhx-2* levels which plays a crucial role in food absorption. RNAi of sodium proton pump that is expressed in the intestine of *C. elegans* delays reproductive senescence and works as a neuroprotectant via decreasing the aggregation of toxic alpha-Synuclein protein *nhx-2* plays a very crucial role in maintaining optimum pH in the gut and facilitates food absorption and also helps in maintaining the food homeostasis. Hence, RNAi of *nhx-2* alters the food absorption thereby mimicking dietary restriction and increasing levels of SIR-2.1 and improving mitochondrial health. We also found that Human estradiol alters the occupancy of epigenetic markers H3K4me3 and H3K9ac on the promoter region of *nhx-2* thereby altering its expression and we anticipate that it corroborates in food absorption via increasing *nhx-2* levels. However, we found that this increase did not alter the aggregation of alpha-Synuclein. We hope to further explore the role of Na+/H+ exchangers as sanative for neurodegeneration. Further, this study is a step towards connecting gut health, estradiol and neuronal health thus encouraging further studies on the gut-brain axis.

## Auther declaration

### Ethics approval and consent to participate

Not applicable

### Consent for publication

Not applicable

### Data Availability Statement

The data that support the findings of this study are available from the corresponding author upon reasonable request.

### Affiliations

Laboratory of Functional Genomics and Molecular Toxicology, Division of Neuroscience and Ageing biology, CSIR-Central Drug Research Institute, Lucknow, UP, 226 031, India Shikha Shukla, Lalit Kumar, Arunabh Sarkar, Kottapalli Srividya and Aamir Nazir

### Contributions

SS conducted the experiments, analysed data, wrote the manuscript, LK, SK and AS contributed in conducting some of the experiments and towards analyzing data; AN conceived the study, provided infrastructure and reagents, analysed the data and edited the manuscript.

### Corresponding author

Correspondence to Aamir Nazir

## Acknowledgments

Authors thank Prof. Tapas K. Kundu and Dr Sweta Sikder for their valuable comments particularly towards the ChIP assay and in editing the manuscript. Strains used in the study were provided by *C. elegans* Genetics Center, University of Minnesottas. SS, LK and AS thank UGC/CSIR for their fellowships. AN acknowledges CSIR for funding support. CDRI Communication number: XXXX

## Research involving Human Participants and/or Animals

Not applicable

## Informed consent

Not applicable

## Supplementary figure

Systematic preliminary screening of genes that delay reproductive senescence in *C. elegans* in regard to various molecular events of neurodegeneration. (1) RNAi of 22 genes available and their effect on □-Syn accumulation: The aggregation of □-Syn was assayed in NL5901 [unc54p::alpha-Synuclein:: YFP+unc-119(+)], a transgenic strain of *C. elegans* that expresses human □-Syn:: YFP transgene in their body wall muscles. All images used the same exposure time and gain settings; Scale bar=100µm. (2) Graphical representation for □-Syn aggregation in transgenic strain NL5901.(3) Effect of RNAi silencing of the various candidates on the relative formation of reactive oxygen (ROS) measured by H2DCFDA assay. H2DCFDA is a chemically reduced, non-fluorescent acetylated form of fluorescein which is readily converted to a green fluorescent form by the activity of ROS. An imbalance between the formation and transmission of ROS has been co-related with PD pathogenesis and can exacerbate its progression. (4) Effect of RNAi silencing of the 22 genes on the number of thrashes in transgenic strain NL5901 of *C. elegans*. The intensity of fluorescence was measured using ImageJ software and Significance was determined using Student□s t-test *p<0.05,**p<0.01,***p<0.001,ns-non-significant.

**Figure S-5:** Interaction between Nuclear Hormonal Receptor and query gene (A)Docking of NHR-14 receptor-estradiol complex with *nhx-2* inverse repeat contig (GAAAAATTGTTCTA,1644-1677,5’-3’).(B) There is no interaction between the NHR-14 and DNA in DNA binding site.

**Figure S-6:** Interaction between Nuclear Hormonal Receptor and query gene (A)Docking of NHR-14 receptor-estradiol complex with her-1 inverse repeat contig (TTTCATATCT,1578-1587,5’-3’).(B) There is an interaction between the NHR-14 complex and DNA contig in DNA binding site. Participating residues are ARG,SER,VAL,TRP,GLN,ASN,LEU,GLU,TYR,THR,PRO,GLY,ILE.

**Figure S-7:** Schematic representation of food absorption

